# Defining a New Standard: Human Platelet Lysate Supports Proliferation and Differentiation of Primary Respiratory Epithelial Cells

**DOI:** 10.64898/2026.08.07.740940

**Authors:** Anna Richter, Jeannine Biermann, Marcus Fulde, Désirée Schaaf

## Abstract

Air-liquid interface (ALI) cultures consisting of well-differentiated primary respiratory epithelial cells (PRECs) provide a versatile *in vitro* model for pharmacological studies and to investigate host-pathogen interactions. Proliferation and differentiation of PRECs require complex media containing several growth factors, hormones, and nutrients. Usually, some of these essential components are provided by the addition of fetal calf serum (FCS). However, several disadvantages of FCS and, most importantly, ethical concerns regarding the method of serum collection have encouraged researchers to find alternatives. Human platelet lysate (hPL) has emerged as a promising alternative to FCS for supporting cell expansion *in vitro*.

In the present study, we investigated the effects of different concentrations of hPL on the proliferation of porcine PRECs and their subsequent differentiation under ALI conditions. Cell morphology was assessed by phase-contrast microscopy, while cell proliferation was evaluated using the ClickTech EdU Cell Proliferation Kit and visualization of proliferating cells by fluorescence microscopy. Differentiation under ALI conditions was monitored by immunofluorescence staining of ciliated cells and the establishment of an intact epithelial barrier was confirmed by measuring transepithelial electrical resistance (TEER).

We found that 5% hPL supported efficient cell growth and the subsequent formation of a functional, well-differentiated airway epithelium comparable to or even better than 10% FCS. Thus, hPL offers a reproducible, ethically sound, and scalable alternative to FCS for complex cell culture models in respiratory research, drug development, and host-pathogen interaction studies.

**Lay Summary:** Respiratory epithelial cells from the lungs of slaughtered animals, such as pigs, can be used for cell culture models to study respiratory diseases and drug development. Air-liquid interface (ALI) cultures closely mimic the natural environment of the airways by exposing the cells to air, making them a valuable alternative to animal experiments. To grow and mature properly, these cells require nutrients and growth factors that are commonly supplied by serum from unborn calves (FCS). However, for ethical and scientific reasons, the use of FCS should be avoided. Therefore, we evaluated whether human platelet lysate (hPL) derived from expired blood donations could replace FCS in ALI cultures. We found that adding 5% hPL to the medium supported efficient cell growth and the development of a well-differentiated airway epithelium.

This approach enables the use of an improved and ethically superior model of the (porcine) respiratory tract in accordance with the 3Rs principle.

## 1. Introduction

The respiratory tract is a vital organ system in both humans and animals, serving essential functions in gas exchange, host defense, and immune regulation. Owing to its central physiological role and its involvement in numerous diseases, the respiratory tract represents a major focus of research across multiple disciplines, including physiology, pharmacology, infectious diseases, immunology, toxicology, and respiratory medicine. A common approach in respiratory research is the use of *in vitro* models that recapitulate key structural and functional features of the respiratory epithelium. These models provide valuable alternatives to animal experimentation and support the implementation of the 3Rs principles (Replacement, Reduction, and Refinement) in biomedical research (Russell and Burch, 1959). Primary respiratory epithelial cells (PRECs) differentiated under air-liquid interface (ALI) conditions are considered one of the most physiologically relevant *in vitro* models of the respiratory epithelium in human respiratory research and are increasingly being adopted in veterinary medicine for the investigation of respiratory physiology, host-pathogen interactions, and therapeutic interventions, which was comprehensively described in a recent review (Weldearegay et al., 2025). ALI cultures recapitulate key structural and functional characteristics of the native airway epithelium, including epithelial polarization, mucociliary differentiation, and barrier integrity (Pezzulo et al., 2011; Prytherch et al., 2011; O’Boyle et al., 2017; Cozens et al., 2018b; Cao et al., 2021). However, their establishment and maintenance are labor-intensive and costly, requiring specialized expertise and careful optimization of culture conditions, particularly medium composition, which critically influences cell proliferation, differentiation, and long-term functionality (Gray et al., 1996; Mao et al., 2009; Cozens et al., 2018a; O’Boyle et al., 2018; Luengen et al., 2020).

Cell proliferation refers to the increase in cell number through mitotic division, whereas differentiation describes the process by which unspecialized cells acquire distinct structural and functional characteristics (Ayers and Jeffery, 1988). Differentiation of PRECs requires a complex combination of supplements, including retinoic acid, insulin, triiodo-L-thyronine, hydrocortisone, transferrin, epinephrin, bovine pituitary extract, and epidermal growth factor (EGF), with concentrations carefully optimized for the species of origin (Gray et al., 1996; Yoon et al., 1997; O’Boyle et al., 2018; Cozens et al., 2018a). Moreover, the composition of the proliferation medium can substantially influence the subsequent differentiation of epithelial cells (O’Boyle et al., 2018; Redman et al., 2024).

Efficient proliferation of PRECs also depends on the presence of several growth-promoting supplements, particularly EGF and Rho-associated protein kinase (ROCK) inhibitors (Dale et al., 2019; Cozens et al., 2018a). In addition, proliferation media are commonly supplemented with serum, most frequently fetal calf serum (FCS; often referred to as fetal bovine serum, FBS), which provides essential growth factors and hormones, transport proteins, minerals and trace elements, lipids, attachment- and spreading factors as well as stabilizing and detoxifying factors (Gstraunthaler, 2003). Despite its widespread use, FCS has several important limitations. As a biologically derived and not precisely defined supplement, it may contain harmful substances such as endotoxins and contaminants (e.g., *Mycoplasma*, viruses) and its composition varies considerably between batches (Price and Gregory, 1982; Barnes et al., 1987; Wessman and Levings, 1999; Stival et al., 2025). Most importantly, the collection of FCS from fetuses obtained from slaughtered pregnant cows has raised significant ethical concerns regarding animal welfare (Jochems et al., 2002; van der Valk et al., 2004). It can be assumed that the fetus has normal brain functions and is therefore capable of feeling pain at the time of blood collection via cardiac puncture (Jochems et al., 2002). Notably, more than two million unborn calves are needed to meet the global demand of approximately 800,000 liters of FCS (Brindley et al., 2012). Consequently, the continued use of FCS as a cell culture supplement for *in vitro* models that were developed to reduce animal experiments seems contradictory and should be abandoned for ethical and scientific reasons.

A promising alternative to FCS as a cell culture substitute is the use of human platelet lysate (hPL) as factors released by activated platelets are known to promote cell attachment, growth, and proliferation (Burnouf et al., 2016). Moreover, hPL is readily available from expired donor platelet concentrates obtained through blood banks and is produced under stringent quality-control procedures, resulting in a highly standardized supplement with reduced batch-to-batch variability. The most simple, efficient, and economic method to induce the release of α-granule components is the physical activation and lysis of platelets by repeated freeze-thaw cycles (Rauch et al., 2011). The capacity of hPL to support epithelial cell culture was demonstrated nearly four decades ago, when it was shown to sustain the differentiation of renal epithelial cells (Gstraunthaler, 1988). Since then, numerous studies have reported the successful use of hPL as a serum substitute for the expansion of several human cell lines and human mesenchymal stromal cells (Burnouf et al., 2016; Barro et al., 2021). However, to the best of our knowledge, the application of hPL for the differentiation of PRECs under ALI conditions has not yet been investigated. Furthermore, studies examining the use of platelet lysates in animal cell culture remain limited. Here, we describe for the first time the successful cultivation of porcine PRECs in proliferation medium supplemented with hPL and evaluate its impact on their subsequent differentiation under ALI conditions.

## Abbreviations

AEGM: Airway Epithelial Cell Growth Medium
ALI: Air-Liquid Interface
DAPI: 4′,6-diamidino-2-phenylindole
EdU: 5-ethynyl-2’-deoxyuridine
EGF: Epidermal Growth Factor
FCS: Fetal Calf Serum
h: hours
hPL: Human Platelet Lysate
IgG: Immunoglobulin
ITCN: Image-based Tool for Counting Nuclei
pPL: Porcine Platelet Lysate
PRECs: Primary Respiratory Epithelial Cells
ROCK: Rho-associated Protein Kinase
RT: Room Temperature
SD: Standard Deviation
TE: Trypsin-EDTA
TEER: Trans-Epithelial Electrical Resistance

## 2. Materials and Methods

A detailed list of reagents used in this study can be found in Tab. S1.

### 2.1. Preparation of Air-Liquid Interface (ALI) Cultures

Porcine respiratory epithelial cells (PRECs) were isolated from the bronchi of freshly slaughtered swine lungs as described in previous publications (Schaaf et al., 2026) (detailed protocol in (Weldearegay et al., 2025)) with some modifications. The lungs came from apparently healthy pigs from regional abattoirs. The separated bronchi were incubated at 4°C for 24 hours (h) in wash medium (Tab. S1) and for 24 h in incubation medium (Tab. S1). PRECs were harvested from the luminal surface of the bronchi and transferred to protease stopping buffer (Tab. S1) to neutralize enzyme activity. To reduce inter-individual variation, cells from at least three different animals were pooled and seeded in collagen I-coated T75 cell culture flasks in Airway Epithelial Cell Growth Medium (AEGM; Tab. S1) supplemented with either 1%/5%/10% heat-inactivated human platelet lysate (hPL) or 10% heat-inactivated fetal calf serum (FCS). Cells were incubated at 37°C and 5% CO_2_ in a humidified atmosphere for up to five days to allow proliferation of the cells (Fig. 1).

**Fig. 1.**
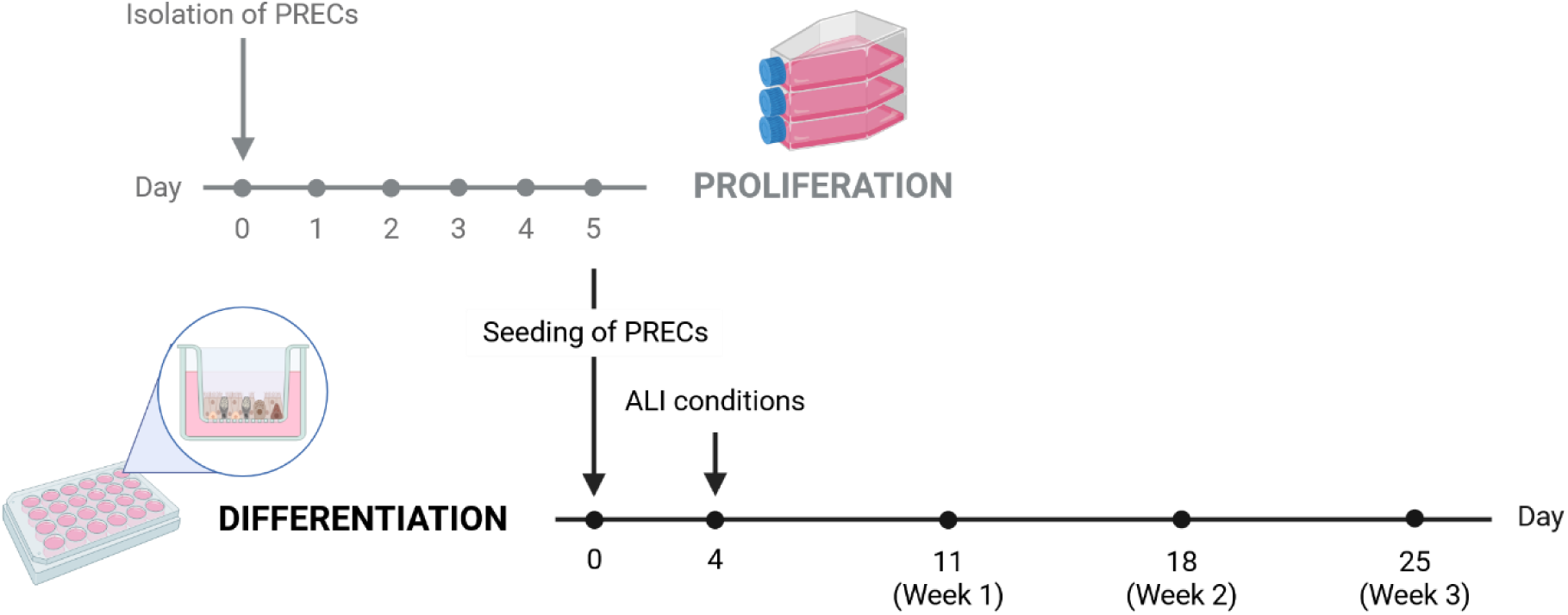
Schematic timeline of porcine respiratory epithelial cell (PREC) proliferation and differentiation. PRECs are isolated from porcine bronchi/trachea and cultivated for five days under submerged conditions in cell culture flasks to allow proliferation. Subsequently, PRECs are dissociated from cell culture flasks and seeded on transwell filters. To facilitate cell proliferation on the filters, PRECs are kept under submerged conditions for four days and then introduced to air-liquid interface (ALI) conditions to allow the development of well-differentiated epithelial cells within the next three weeks. Created in BioRender. Schaaf, D. (2026) https://BioRender.com/turm94w.

When cells have grown confluent, they were detached using 0.05% trypsin-EDTA (TE) and seeded on collagen IV-coated cell culture inserts (polycarbonate membrane, 6.5 mm diameter, 0.4 µm pore size; VWR, Cat. No. 734-2742; 2.5 × 10^5^ cells/insert). During the next four days, PRECs were cultivated under submerged conditions in AEGM supplemented with either hPL or FCS to allow the formation of a confluent cell layer. Subsequently, medium from the apical compartment was removed to expose PRECs to air and the medium in the basal compartment was replaced by ALI medium (Tab. S1), which contains neither hPL nor FCS. Under these so called “ALI conditions”, PRECs can differentiate and build a pseudostratified epithelial barrier consisting of ciliated and mucus-producing cells within 2-3 weeks (Fig. 1). ALI cultures were incubated at 37°C and 5% CO_2_ in a humidified atmosphere, supplied with fresh medium every 2-3 days and washed once a week to remove excess mucus and dead cells.

Integrity of the epithelial barrier was determined by measurement of the transepithelial electrical resistance (TEER) using a volt-ohm meter (EVOM™ Manual equipped with a STX4 electrode, World Precision Instruments). At least two independent TEER measurements were performed with technical duplicates each.

### 2.2. Phase Contrast and Immunofluorescence Microscopy

Cell morphology of freshly isolated and proliferating PRECs in AEGM supplemented with either 1%, 5%, or 10% human platelet lysate (hPL) or 10% fetal calf serum (FCS) was monitored by phase contrast microscopy. Representative images of randomly selected fields of view were acquired with the Keyence BZ-X800E, an inverted fluorescence phase contrast microscope, equipped with the Keyence Plan Fluorite 20X/0.45 LD PH air objective lens. Brightness and contrast were adjusted using BZ-X800 Analyzer software (version 1.1.2.4, Keyence).

The extent of cilia formation during differentiation of PRECs under ALI conditions was visualized by immunofluorescence staining as previously described (Schaaf et al., 2026). PRECs were fixed with 3.7% formaldehyde and then incubated for at least 1 h in blocking buffer at room temperature (RT). The monoclonal CY3-conjugated anti-β-tubulin antibody was diluted 1:500 in antibody dilution buffer and incubated overnight at 4°C to stain cilia. Nuclei were colored with 4′,6-diamidino-2-phenylindole (DAPI). Afterwards, the membrane was removed from the insert and embedded in ProLong Gold Antifade Reagent. ALI cultures were analyzed with the Keyence BZ-X800E. Composite images of the entire membrane were generated from individual tiles captured with the Plan Apochromat 10X/0.45 air objective lens using the stitching acquisition mode and processed with the BZ-X800 Analyzer software. Image stacks with a z-distance of 0.3 µm were captured with the Plan Apochromat 40X/0.95 air objective lens and subsequently merged using the BZ-X800 Analyzer software. The software was also used to adjust colors as well as brightness and contrast.

### 2.3. Proliferation Assay

Proliferation capacity of PRECs was assessed using the ClickTech EdU Cell Proliferation Kit 488 Sensitive for Imaging (Carl Roth, Cat. No. 1Y57.1) according to the manufacturer’s instructions. PRECs were isolated from porcine bronchi as described above and cultivated in collagen I-coated T75 cell culture flasks in AEGM supplemented with 5% hPL until confluence (approximately five days). Then, cells were detached using 0.05% TE, cryo-preserved in freezing medium (Tab. S1; 5 × 10^6^ cells/vial), and stored in liquid nitrogen. Frozen PRECs were thawed and seeded at a density of 2 × 10^5^ cells/well on collagen I-coated glass cover slips in a 24-well plate in AEGM supplemented with either 1%/5%/10% heat-inactivated hPL or 10% heat-inactivated FCS. Cells were incubated at 37°C and 5% CO_2_ in a humidified atmosphere for 24 h. The next day, PRECs were labeled with the thymidine analog EdU (5-ethynyl-2’-deoxyuridine) by adding 5 µM EdU to AEGM supplemented with either 1%/5%/10% hPL or 10% FCS and incubated for up to three days (Fig. 2). Every day, samples were collected for fixation with 4% formaldehyde and permeabilization with 1% saponin. Subsequently, the click chemistry reaction cocktail was added and incubated for 30 minutes (min) at RT. Finally, nuclei were stained with DAPI and the cover slips were embedded in ProLong Gold Antifade Reagent. Image acquisition was performed with the Keyence BZ-X800E, equipped with the Plan Apochromat 10X/0.45 air objective lens, using the stitching acquisition mode. Individual image tiles were assembled into composite images using the BZ-X800 Analyzer software. Additionally, colors as well as brightness and contrast were adjusted with this software.

**Fig. 2.**
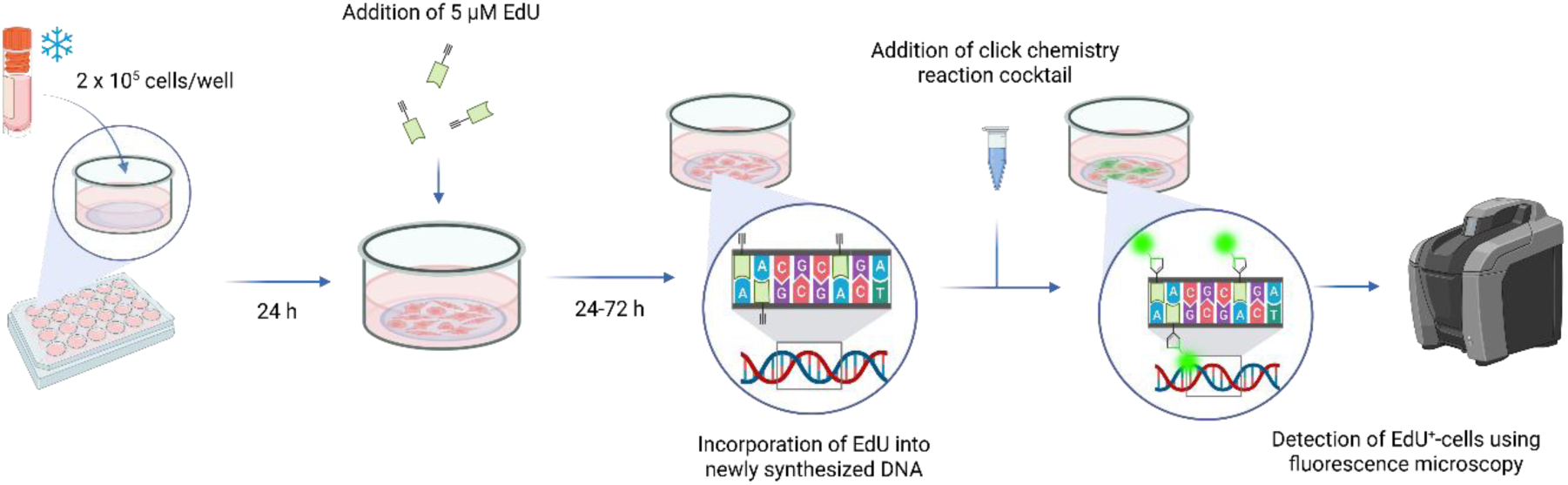
Schematic presentation of the proliferation assay. Proliferating PRECs were labeled with the ClickTech Sensitive EdU Cell Proliferation Kit for Imaging and detected by fluorescence microscopy. Created in BioRender. Schaaf, D. (2026) https://BioRender.com/vw2kmwr.

### 2.4. Nuclei Counting

As described above, proliferating PRECs were labeled using the ClickTech Sensitive EdU Cell Proliferation Kit for Imaging (EdU^+^ cells; Fig. 2) and nuclei of all cells were stained with DAPI. Individual images acquired with the Keyence BZ-X800E, equipped with the Plan Apochromat 10X/0.45 air objective lens, (see 2.3) were analyzed using the ImageJ plug-in Image-based Tool for Counting Nuclei (ITCN; https://github.com/PMB-KU/CountNuclei). For this, individual, single-color images were pre-processed by converting the image to 8-bit and inverting bright and dark signals using ImageJ software (version 1.54g; National Institutes of Health, USA). Brightness and contrast were automatically adjusted by the software. Subsequently, pre-processed images were automatically analyzed using Batch ITCN (Edge: 0 pixels; Nuclei diameter: 30 pixels; Minimum distance: 15 pixels; Threshold: 0.2 for DAPI and 0.1 for EdU^+^ cells, respectively) counting dark signals on bright background (Fig. S1). The number of EdU^+^ cells was expressed as proportion (%) of proliferating cells from total cell count (DAPI-labeled nuclei). Images containing air bubbles, fibrous particles, and other artifacts were excluded from the analysis. At least 46 images/sample (on average 58 images/sample, 1-2 technical replicates each) of the randomly selected stitching area were analyzed, and three independent experiments were performed. Fig. 4B and 4C show the average value of all analyzed images per sample and per experiment for total cell numbers and % EdU^+^ cells, respectively.

Nuclei counting for ALI cultures was performed as described for proliferating PRECs with some modifications. Image acquisition was performed with the Keyence BZ-X800E, equipped with the Plan Apochromat 20X/0.75 air objective lens, using the stitching acquisition mode. Individual images of nuclei stained with DAPI were pre-processed as described above and automatically analyzed using Batch ITCN (Edge: 0 pixels; Nuclei diameter: 20 pixels; Minimum distance: 10 pixels; Threshold: 0.1) (Fig. S2). Images containing any artefacts interfering with nuclei counts were excluded from the analysis. At least 35 images/filter (on average 56 images/filter, two technical replicates) were analyzed, and three independent experiments were performed. Fig. 4A shows the average value of all analyzed images per sample and per experiment, while Fig. S6 shows all nuclei counts per image from two technical replicates for each independent experiment.

### 2.5. Statistical Analysis

Unless otherwise specified in the figure caption, data are shown as medians and individual values from at least three independent experiments (in technical duplicates). For reasons of clarity, TEER values are shown as mean ± standard deviation (SD) from 2-3 independent experiments (in technical duplicates). All statistical analyses were carried out using GraphPad Prism version 10.6.1 for Windows (GraphPad Software). Statistical significance among several groups was analyzed using the Kruskal-Wallis test followed by Dunn’s multiple comparisons test for small sample sizes (n = 3) or by one-way ANOVA followed by Tukey’s multiple comparisons test when normal distribution of the data could be assumed (Fig. S6). Statistical significance between two groups was analyzed using Mann-Whitney test (Fig. S5). *P* < 0.05 was considered significant.

## 3. Results

### 3.1. Proliferation of primary porcine respiratory epithelial cell (PRECs)

After isolation of PRECs from porcine bronchi, the cells were cultivated under submerged conditions (covered with cell culture medium) in cell culture flasks to allow cell proliferation until confluence. During proliferation of PRECs in Airway Epithelial Cell Growth Medium (AEGM) supplemented with either 1%/5%/10% human platelet lysate (hPL) or 10% fetal calf serum (FCS), we monitored cell morphology by light microscopy (Fig. 3 and Fig. S3). Interestingly, we found considerably less cells when AEGM was supplemented with 1% hPL only, whereas cell numbers appeared comparable when AEGM was supplemented with 5% hPL, 10% hPL, or 10% FCS. All cells showed the typical epithelial polygonal or cobblestone morphology, which was especially apparent during the first two days when cells were not yet confluent. Independent of the cultivation medium, we noticed a granular cytoplasm in all cells and vacuolization in some PRECs (Fig. 3 and Fig. S3).

**Fig. 3.**
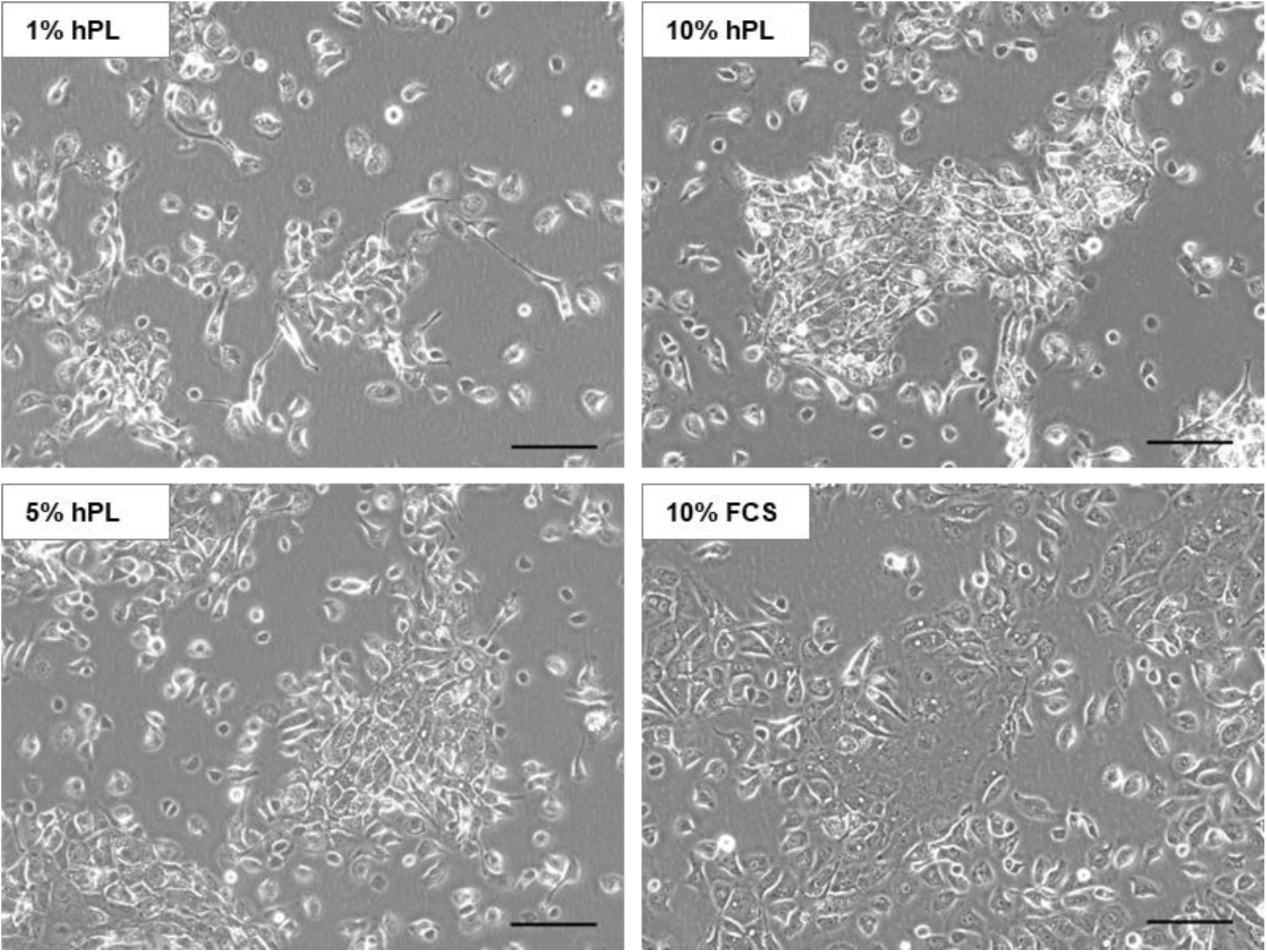
Cell morphology of proliferating PRECs. Freshly isolated PRECs were cultivated in Airway Epithelial Cell Growth Medium (AEGM) supplemented with either 1%, 5%, or 10% human platelet lysate (hPL) or 10% fetal calf serum (FCS) for up to five days. Cell morphology was monitored by phase contrast microscopy. Representative images of cells cultivated for two days under submerged conditions are shown. Bars represent 100 µm.

The number of proliferating cells was determined using the ClickTech Sensitive EdU Cell Proliferation Kit for Imaging. The thymidine analog EdU (5-ethynyl-2’-deoxyuridine) is incorporated into DNA during its synthesis and fluorescently labeled nuclei of proliferating cells can be detected by fluorescence microscopy (Fig. 4A and Fig. S4). We counted the total number of cells (labeled with 4′,6-diamidine-2′-phenylindole/DAPI) as well as the number of proliferating (EdU-positive/EdU^+^) cells after 2-4 days of cultivation in AEGM supplemented with hPL or FCS, respectively. Interestingly, in contrast to the results described above for the freshly isolated PRECs, the total cell count was highest when AEGM was supplemented with 1% or 5% hPL - it increased during the first two days and slightly decreased afterwards-, whereas cell counts were lower when PRECs were cultured in AEGM supplemented with 10% hPL or 10% FCS (Fig. 4B). Notably, after four days of cultivation, the total cell number was lower in the presence of EdU than in the absence of EdU, which was particularly prominent when PRECs were cultivated in AEGM supplemented with 10% hPL or 10% FCS (Fig. S5). During the first days of cultivation, the percentage of proliferating cells (number of EdU^+^ cells compared to the total cell number) was highest in PRECs cultivated in AEGM supplemented with 10% FCS and 20-30% lower in cells cultivated in AEGM supplemented with hPL independent of its concentration. However, four days after seeding, the proportion of proliferating cells was lowest in PRECs cultivated in AEGM supplemented with 5% hPL and comparable in cells cultivated in AEGM supplemented with 1% hPL, 10% hPL, or 10% FCS (Fig. 4C).

**Fig. 4.**
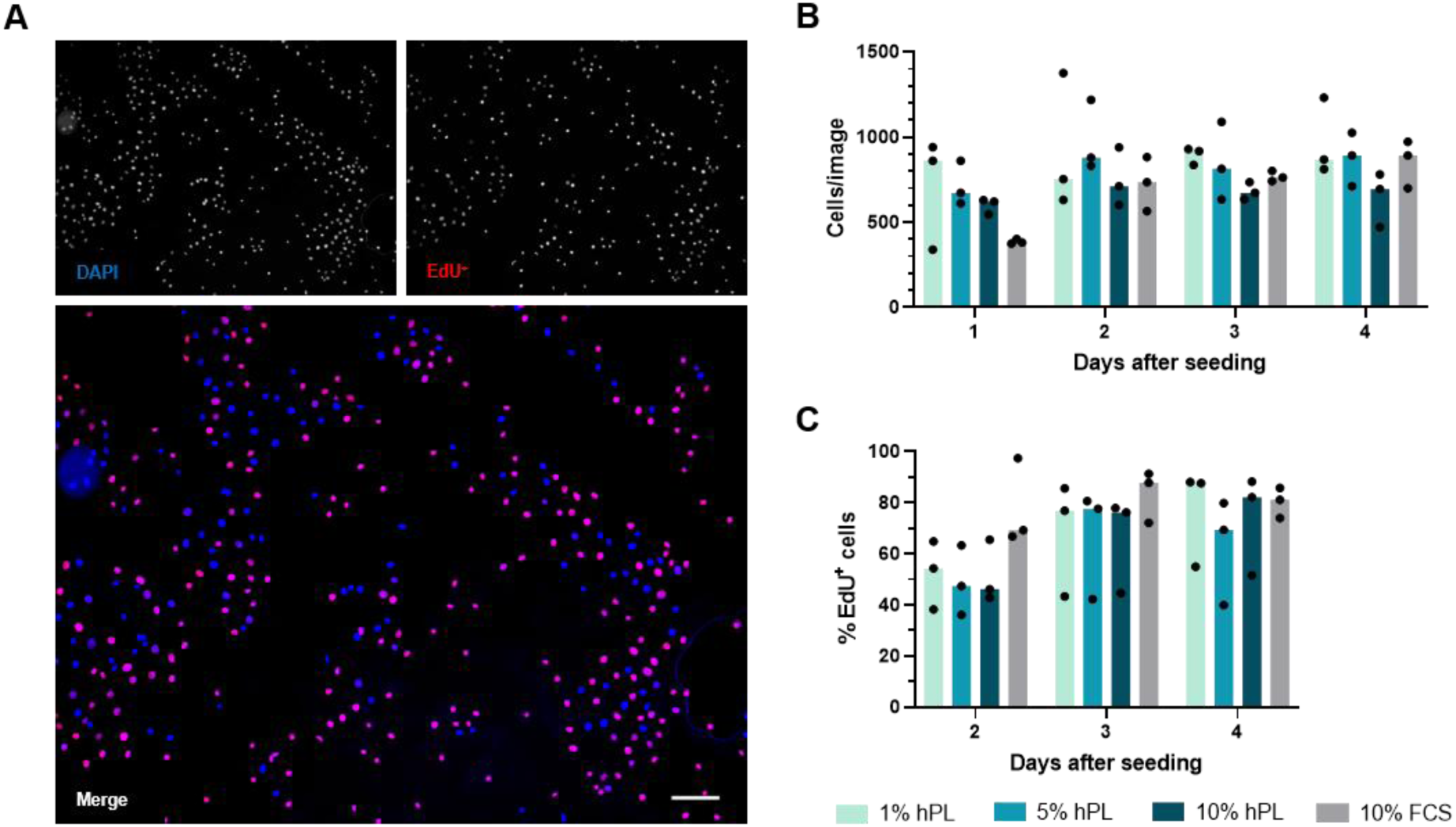
Proliferation capacity of PRECs. Frozen PRECs were seeded at a density of 2 × 10^5^ cells/well on glass cover slips in a 24-well plate. AEGM was supplemented with either 1%, 5%, or 10% hPL or 10% FCS. After one day, medium was changed and 5 µM 5-ethynyl-2’-deoxyuridine (EdU) were added to the cells to label proliferating PRECs. Proliferating (EdU^+^) cells were then detected by fluorescence microscopy. (A) Representative image of nuclei labeled with 4′,6-diamidine-2′-phenylindole (DAPI; blue) and proliferating cells labeled with EdU (red) two days after seeding. Bar represents 100 µm. (B) The number of cells per image was determined by counting nuclei stained with DAPI using the ImageJ plug-in Image-based Tool for Counting Nuclei (ITCN) in a randomly selected area. (C) The number of EdU^+^ cells was counted using the ImageJ plug-in ITCN and expressed as proportion (%) of proliferating cells from total cell number. The median of three independent experiments is shown. Significant differences between hPL and 10% FCS were analyzed with Kruskal-Wallis test followed by Dunn’s multiple comparisons test (*P* < 0.05 was considered significant).

In summary, these results show that PRECs cultivated in AEGM supplemented with hPL better attached initially, while more PRECs proliferated when cultivated in AEGM supplemented with 10% FCS. However, the overall cell number and cell morphology of PRECs in 5% hPL is comparable to PRECs in 10% FCS for both freshly isolated and frozen PRECs.

### 3.2. Differentiation of PRECs under air-liquid interface (ALI) conditions

Previous studies have shown that composition of the proliferation medium has a considerable impact on differentiation of epithelial cells (O’Boyle et al., 2018; Redman et al., 2024). Therefore, we evaluated the differentiation potential of PRECs that had previously been cultivated in AEGM supplemented with either hPL or FCS in cell culture flasks. PRECs were seeded on transwell filters and cultivated under ALI conditions, which facilitates the differentiation into a pseudostratified epithelium consisting of mucus-producing and ciliated epithelial cells (Whitcutt et al., 1988). Before introducing the cells to ALI conditions, PRECs were cultivated for four days under submerged conditions in AEGM supplemented with either 1%/5%/10% hPL or 10% FCS to allow proliferation of the cells on the filter membrane. Starting on the fourth day, all cells were cultivated with the same medium (ALI medium) in the basal compartment, which does not contain any hPL or FCS, and the apical compartment was exposed to air. We determined cell numbers, transepithelial electrical resistance (TEER), and the development of cilia every week starting from the beginning of ALI conditions. Independent of the media conditions, we counted approximately 5 × 10^5^ cells/membrane by fluorescence microscopy throughout the observation period (Fig. 5A).

**Fig. 5.**
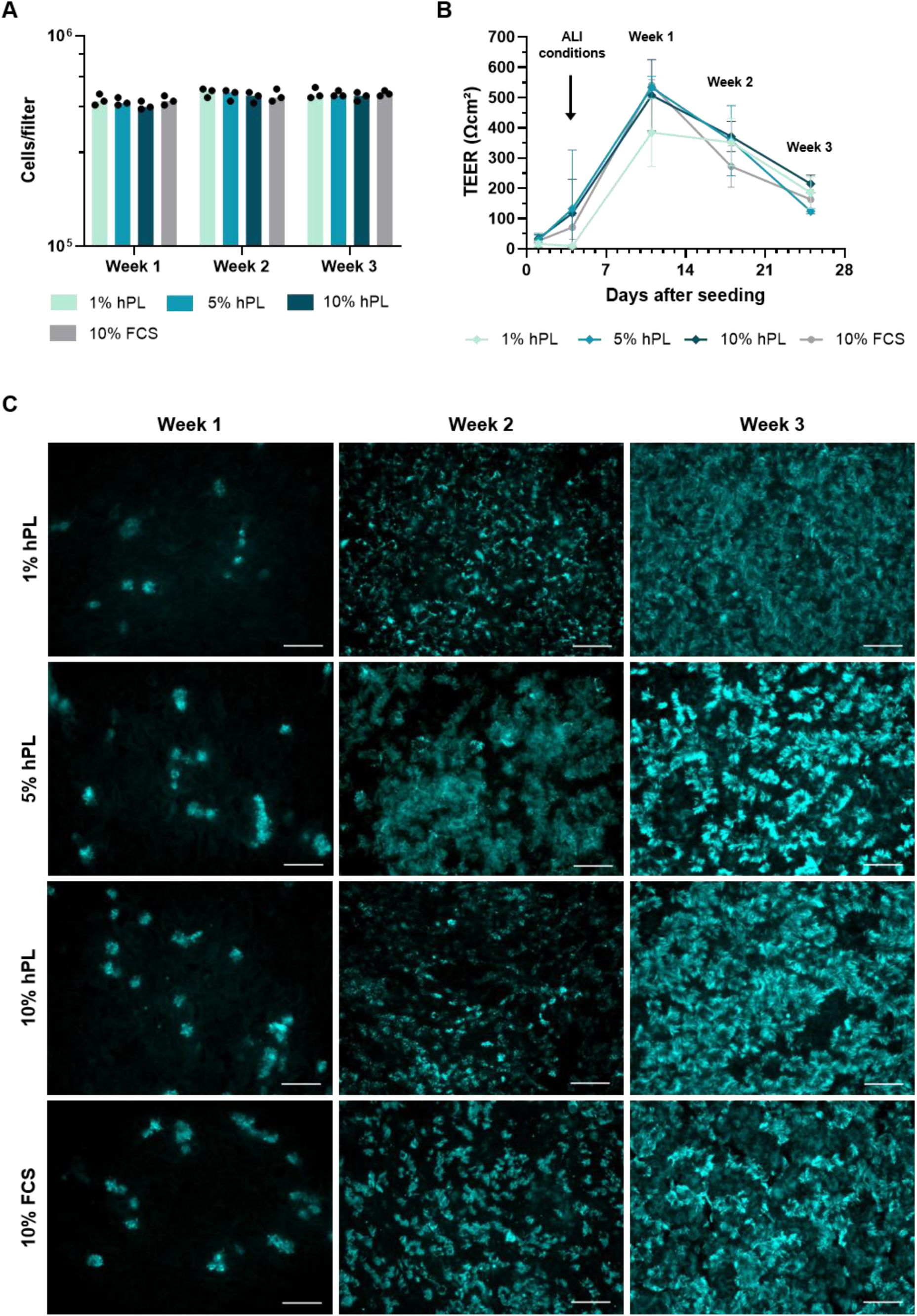
Differentiation of PRECs under air-liquid interface (ALI) conditions. Before PRECs were exposed to ALI conditions, they have been cultivated on transwell filters under submerged conditions in AEGM supplemented with either 1%, 5%, or 10% hPL or 10% FCS for four days. Subsequently, PRECs were differentiated under ALI conditions in medium without any hPL or FCS for up to three weeks. (A) The number of cells/filter was calculated by nuclei counting using the ImageJ plug-in ITCN. After labeling with DAPI, nuclei of at least 35 images/filter were automatically counted and the number of cells/image was extrapolated to the number of cells/filter. The median of three independent experiments is shown (two replicates each). Significant differences between hPL and 10% FCS were analyzed with Kruskal-Wallis test followed by Dunn’s multiple comparisons test. (B) Epithelial barrier integrity was evaluated by trans-epithelial electrical resistance (TEER) measurement and expressed as resistance relative to the surface area (Ωcm^2^). Mean ± SD of 2-3 independent experiments is shown. Significant differences between hPL and 10% FCS were analyzed with Kruskal-Wallis test followed by Dunn’s multiple comparisons test. (C) Immunofluorescence staining of cilia (β-tubulin) as an indicator of the level of differentiation under ALI conditions. Bars represent 50 µm.

However, when comparing the nuclei counts/image for all replicates individually, we noticed a slight increase in nuclei counts from week 1 to week 2 (Fig. S6). Notably, the lowest cell numbers were counted when PRECs were cultivated in AEGM supplemented with 10% hPL and highest cell numbers with 1% hPL, whereas cultivation in AEGM supplemented with 5% hPL and 10% FCS was almost comparable (Fig. S6). Measurement of TEER revealed a typical course peaking after one week under ALI conditions and a subsequent decline until the values stabilized. TEER values were comparable for 5% hPL, 10% hPL, and 10% FCS during the first week but slightly lower for cells cultivated in AEGM supplemented with 10% FCS after two weeks under ALI conditions. Interestingly, cells grown in 1% hPL showed lower TEER values during the first 11 days on transwell filters but comparable values to the other hPL concentrations during the remaining observation period (Fig. 5B). A common method to monitor differentiation of PRECs under ALI conditions is the visualization of ciliated epithelial cells by immunofluorescence microscopy. For all media conditions, we noticed an increase of ciliated cells during the observation period. Already after two weeks under ALI conditions, PRECs cultivated in AEGM supplemented with 5% hPL showed strong ciliation, which was even higher compared to cells grown in 10% FCS (Fig. 5C and Fig. S7).

Finally, immunofluorescence staining, nuclei counting, and TEER measurement revealed that the initial cultivation of PRECs with 5% hPL can efficiently support their subsequent differentiation under ALI conditions.

## 4. Discussion

A recent review (Weber et al., 2025) comprehensively summarizes the pitfalls associated with fetal calf serum (FCS), which is a complex and poorly defined mixture of growth factors, hormones, amino acids, carbohydrates, lipids, and other biologically active components (Zheng et al., 2006; van der Valk et al., 2018). Although FCS efficiently supports cell growth and proliferation, there is evidence indicating that FCS might exert undesirable effects on cell physiology (Khodabukus and Baar, 2014; Kwon et al., 2016; van Vijven et al., 2021). Moreover, substantial batch-to-batch variability in FCS composition, together with the risk of contamination by endotoxins or pathogens, requires extensive batch testing, increases experimental costs, and compromises reproducibility (Baker, 2016; Zhang et al., 2022; Liu et al., 2023; Stival et al., 2025). In particular, the collection of serum from bovine fetuses via cardiac puncture raises serious ethical concerns regarding animal welfare as the animals can be expected to be conscious and suffering from pain (Jochems et al., 2002; van der Valk et al., 2004). Consequently, alternatives are urgently needed but literature on FCS alternatives in animal cell culture as well as in complex cell culture models, such as air-liquid interface (ALI) cultures is sparse. In a recent study, cell culture medium for porcine ALI cultures was supplemented with an FCS-equivalent, which is derived from bovine animals after birth promising a higher lot-to-lot consistency, stable pricing, and superior traceability (Castillo-Espinoza et al., 2026) (https://www.atlasbio.com/ab-product/equalfetal/; last access 2026-07-17). Nevertheless, many of the disadvantages described for FCS might still apply for this FCS-equivalent. Thus, chemically defined media supplemented with recombinant proteins represent a more transparent, reproducible, and ethically acceptable alternative that can minimize contamination risks while improving experimental consistency (Gstraunthaler, 2003; van der Valk et al., 2010). However, developing optimal media formulation is challenging and time-consuming, as media requirements are highly cell type specific. Furthermore, commercially available chemically defined media are still unavailable for many cell types, particularly for animal cell cultures.

Platelets play an important role in tissue regeneration and contain substantial amount of different growth factors that are released upon activation (Golebiewska and Poole, 2015; Burnouf et al., 2016). Four decades ago, human platelet lysate (hPL) was introduced as a cell culture supplement efficiently supporting proliferation of human tumor cell lines, endothelial cells, and fibroblast (Hara et al., 1980; King and Buchwald, 1984; Gstraunthaler, 1988; Umeno et al., 1989). Since then, research on hPL has primarily focused on its application in human stem cell culture (Burnouf et al., 2016; Guiotto et al., 2020), whereas studies investigating its effects on epithelial cells remain limited. To our knowledge, hPL has not yet been evaluated as a supplement for primary animal-derived epithelial cells or for the differentiation of primary respiratory epithelial cells (PRECs) under ALI conditions. Hence, we have tested the suitability of hPL to support proliferation of porcine PRECs and their subsequent differentiation under ALI conditions. PRECs cultivated in Airway Epithelial Cell Growth Medium (AEGM) supplemented with 5% hPL proliferated successfully and built a well-differentiated respiratory epithelium under ALI conditions characterized by a dense carpet of cilia and high epithelial barrier integrity. Even the lowest concentration of hPL was sufficient to support proliferation and differentiation of PRECs, although the development of a stable epithelial barrier was a little delayed. Further investigations are needed in the future to evaluate whether cultivation of PRECs with hPL has an impact on the ciliary beat frequency (Roth et al., 2025), on the cell composition (i.e., the presence and the ratio of basal cells, goblet/secretory cells, ciliated cells and other specialized cells), or on the transcriptional profile of the cells (Pezzulo et al., 2011).

However, despite its advantages, hPL has several important limitations that should be considered. Growth factor composition varies between donors, platelet preparation and activation methods, and pooling strategies, which is due to the lack of standardization in manufacturing (Schallmoser and Strunk, 2013; Burnouf et al., 2016). This batch-to-batch variability can lead to differences in cell proliferation and phenotype, but these negative effects can be leveled out by pooling platelets from several different donors and is usually the case for commercially available hPL (Burnouf et al., 2016; Bieback et al., 2019). Next, it has to be taken into consideration that hPL preparations contain fibrinogen forming clots when mixed with calcium-containing media (Schallmoser and Strunk, 2013; Burnouf et al., 2016), requiring the addition of anticoagulants, such as heparin, or the depletion of fibrinogen. In our study, we used fibrinogen-depleted hPL, which is commercially available, to avoid the addition of heparin and the laborious adjustment of its optimal concentration. Compared to FCS, hPL contains significantly higher amounts of immunoglobulins (IgG) (Duarte Rojas et al., 2024), which is generally not considered a major limitation. However, when performing immunological and proteomic studies, the high content of IgG should be taken into account as the high protein concentration can mask low abundant proteins (Tu et al., 2010) and IgG can modulate immune cell behavior by engaging their Fc-receptor (Nimmerjahn and Ravetch, 2008) or may interfere with immunological assays, particularly antibody-based detection assays (Tate and Ward, 2004). Moreover, Jan *et al*. have suggested that the presence of biologically active factors in hPL, such as cytokines, may be responsible for the induction of an antiviral inflammatory response in hPL-cultured glioblastoma cells (Jan et al., 2022). Heat-inactivation at 56°C for 30 min can be performed to inactivate complement components, but it is important to note that heat-labile growth factors can also be inactivated by this process, whereas it is not sufficient to reduce the concentration or biological presence of immunoglobulins considerably (Anitua et al., 2014).

Instead of using hPL for porcine PRECs, it seems more reasonable to use porcine platelet lysates (pPL) as substitute for FCS to follow a xeno-free approach. Indeed, other researchers have used pPL successfully as a supplement for animal cell culture (Vero cells and Chinese Hamster Ovary cells) and reported low batch-to-batch variability (Alden et al., 2007). However, to our knowledge, pPL was not yet tested as a supplement for primary porcine cell cultures, especially not for cultures as complex as ALI cultures. Moreover, production of pPL must first be established in the laboratory as it is not commercially available. Since the composition of hPL and pPL is comparable (Alden et al., 2007; Duarte Rojas et al., 2024), we assume that the cultivation of PRECs using pPL would also be successful and that it would therefore make sense to establish pPL production in the future. Another research group reported the use of porcine serum instead of FCS in human and porcine stem cell culture. Both cell types, human adipose-derived mesenchymal stem/stromal cells and porcine skeletal muscle–derived satellite cells, were able to proliferate and maintained their differentiation capacity (Hahn et al., 2024). This approach seems very promising, above all because porcine serum can be obtained easily and cost-effectively from pig blood as a side-product of slaughtering being a sustainable and readily available source (Hahn et al., 2024). Future studies will tell if porcine serum is also suitable as a supplement for cultivation of porcine PRECs or other porcine cells.

FCS can protect cells during freezing because of its high protein concentration, which is why medium for cryopreservation of cells is usually supplemented with 40% FCS. In the present study, we also followed this protocol, in part because the use of hPL (or pPL) as a cryoprotective agent is not yet well described and needs to be established for porcine PRECs. Rojas *et al*. compared the efficiency of FCS, hPL, and hPL serum in cryopreservation of human dermal fibroblasts and human mesenchymal stem cells, wherein the freezing medium consisted of 90% hPL/hPL serum/FCS and 10% dimethyl sulfoxide (DMSO). The authors stated that hPL serum was a better replacement for FCS than hPL with regard to cell viability and proliferation post-thawing indicating that hPL might not be an effective cryoprotective agent (Duarte Rojas et al., 2024). Other promising approaches for FCS-free cryopreservation of porcine PRECs, such as the use of commercially available FCS-free cryopreservation media, the enrichment of the freezing medium with high protein concentrations (e.g. by adding bovine serum albumin), or the sole addition of DMSO to the freezing medium (Weber et al., 2025), will be evaluated in the future.

To our knowledge, this study is the first to demonstrate that hPL can successfully replace FCS for the expansion of porcine PRECs and their subsequent differentiation under ALI conditions. These findings confirm hPL as a promising alternative for complex respiratory epithelial culture systems and represent an important step toward reducing the use of FCS in accordance with the 3Rs principles. While further optimization, including the development of species-specific supplements such as pPL and fully FCS-free cryopreservation, is still required, our work provides a foundation for more ethical, reproducible, and physiologically relevant *in vitro* models.

## Acknowledgements

This work was financially supported by the Deutsche Forschungsgemeinschaft (DFG; SCHA 2406/1-1), AR was additionally supported by the Hannover Graduate School for Neurosciences, Infection Medicine and Veterinary Sciences (HGNI) of the University of Veterinary Medicine Hannover, and MF received funding by the DFG through the Research Training Group GRK 3051 (Project-ID 529208971). The authors thank the slaughterhouses Twachtmann (Nienburg, Germany) and Klos (Nienhagen, Germany) for providing swine lungs. Furthermore, the authors acknowledge consultation from Silke Isenhardt (PL Bioscience, Aachen, Germany) about hPL and from Simone Bergmann (Technical University Braunschweig, Germany) about the proliferation assay.

## Data Availability

The main research data supporting the findings of this study are included in this manuscript and its supplementary materials. The raw data are available from the corresponding author upon reasonable request.

## Author Contributions

AR: Formal Analysis, Investigation, Visualization, Writing – Review C Editing

JB: Investigation, Writing – Review C Editing

MF: Conceptualization, Writing – Review C Editing

DS: Conceptualization, Formal Analysis, Funding Acquisition, Methodology, Project Administration, Supervision, Validation, Visualization, Writing – Original Draft Preparation, Writing – Review C Editing

## References

Alden, A., Gonzalez, L., Persson, A. et al. (2007). Porcine platelet lysate as a supplement for animal cell culture. Cytotechnology 55, 3–8. doi:10.1007/s10616-007-9097-9

Anitua, E., Muruzabal, F., De la Fuente, M. et al. (2014). Effects of heat-treatment on plasma rich in growth factors-derived autologous eye drop. Exp Eye Res 119, 27–34. doi:10.1016/j.exer.2013.12.005

Ayers, M. M. and Jeffery, P. K. (1988). Proliferation and differentiation in mammalian airway epithelium. Eur Respir J 1, 58–80.

Baker, M. (2016). Reproducibility: Respect your cells! Nature 537, 433–435. doi:10.1038/537433a

Barnes, D., McKeehan, W. L. and Sato, G. H. (1987). Cellular endocrinology: Integrated physiology in vitro. In Vitro Cell Dev Biol 23, 659–662. doi:10.1007/BF02620978

Barro, L., Burnouf, P. A., Chou, M. L. et al. (2021). Human platelet lysates for human cell propagation. Platelets 32, 152–162. doi:10.1080/09537104.2020.1849602

Bieback, K., Fernandez-Munoz, B., Pati, S. et al. (2019). Gaps in the knowledge of human platelet lysate as a cell culture supplement for cell therapy: A joint publication from the aabb and the international society for cell & gene therapy. Cytotherapy 21, 911–924. doi:10.1016/j.jcyt.2019.06.006

Brindley, D. A., Davie, N. L., Culme-Seymour, E. J. et al. (2012). Peak serum: Implications of serum supply for cell therapy manufacturing. Regen Med 7, 7–13. doi:10.2217/rme.11.112

Burnouf, T., Strunk, D., Koh, M. B. et al. (2016). Human platelet lysate: Replacing fetal bovine serum as a gold standard for human cell propagation? Biomaterials 76, 371–387. doi:10.1016/j.biomaterials.2015.10.065

Cao, X., Coyle, J. P., Xiong, R. et al. (2021). Invited review: Human air-liquid-interface organotypic airway tissue models derived from primary tracheobronchial epithelial cells-overview and perspectives. In Vitro Cell Dev Biol Anim 57, 104–132. doi:10.1007/s11626-020-00517-7

Castillo-Espinoza, A. F., Nelli, R. K., Mora-Diaz, J. C. et al. (2026). Mycoplasma hyopneumoniae modulates ciliary function and epithelial integrity in air-liquid interface porcine respiratory epithelial cells (ali-precs). Microbiol Spectr e0199825. doi:10.1128/spectrum.01998-25

Cozens, D., Grahame, E., Sutherland, E. et al. (2018a). Development and optimization of a differentiated airway epithelial cell model of the bovine respiratory tract. Sci Rep 8, 853. doi:10.1038/s41598-017-19079-y

Cozens, D., Sutherland, E., Marchesi, F. et al. (2018b). Temporal differentiation of bovine airway epithelial cells grown at an air-liquid interface. Sci Rep 8, 14893. doi:10.1038/s41598-018-33180-w

Dale, T. P., Borg D’anastasi, E., Haris, M. et al. (2019). Rock inhibitor y-27632 enables feeder-free, unlimited expansion of sus scrofa domesticus swine airway stem cells to facilitate respiratory research. Stem Cells Int 2019, 3010656. doi:10.1155/2019/3010656

Duarte Rojas, J. M., Restrepo Munera, L. M. and Estrada Mira, S. (2024). Comparison between platelet lysate, platelet lysate serum, and fetal bovine serum as supplements for cell culture, expansion, and cryopreservation. Biomedicines 12, doi:10.3390/biomedicines12010140

Golebiewska, E. M. and Poole, A. W. (2015). Platelet secretion: From haemostasis to wound healing and beyond. Blood Rev 29, 153–162. doi:10.1016/j.blre.2014.10.003

Gray, T. E., Guzman, K., Davis, C. W. et al. (1996). Mucociliary differentiation of serially passaged normal human tracheobronchial epithelial cells. Am J Respir Cell Mol Biol 14, 104–112. doi:10.1165/ajrcmb.14.1.8534481

Gstraunthaler, G. (2003). Alternatives to the use of fetal bovine serum: Serum-free cell culture. ALTEX 20, 275–281.

Gstraunthaler, G. J. (1988). Epithelial cells in tissue culture. Ren Physiol Biochem 11, 1–42. doi:10.1159/000173147

Guiotto, M., Raffoul, W., Hart, A. M. et al. (2020). Human platelet lysate to substitute fetal bovine serum in hmsc expansion for translational applications: A systematic review. J Transl Med 18, 351. doi:10.1186/s12967-020-02489-4

Hahn, O., Peters, K., Hartmann, A. et al. (2024). Potential of animal-welfare compliant and sustainably sourced serum from pig slaughter blood. Cell Tissue Res 397, 205–214. doi:10.1007/s00441-024-03904-8

Hara, Y., Steiner, M. and Baldini, M. G. (1980). Platelets as a source of growth-promoting factor(s) for tumor cells. Cancer Res 40, 1212–1216.

Jan, M. W., Chiu, C. Y., Chen, J. J. et al. (2022). Human platelet lysate induces antiviral responses against parechovirus a3. Viruses 14, doi:10.3390/v14071499

Jochems, C. E., van der Valk, J. B., Stafleu, F. R. et al. (2002). The use of fetal bovine serum: Ethical or scientific problem? Altern Lab Anim 30, 219–227. doi:10.1177/026119290203000208

Khodabukus, A. and Baar, K. (2014). The effect of serum origin on tissue engineered skeletal muscle function. J Cell Biochem 115, 2198–2207. doi:10.1002/jcb.24938

King, G. L. and Buchwald, S. (1984). Characterization and partial purification of an endothelial cell growth factor from human platelets. J Clin Invest 73, 392–396. doi:10.1172/JCI111224

Kwon, D., Kim, J. S., Cha, B. H. et al. (2016). The effect of fetal bovine serum (fbs) on efficacy of cellular reprogramming for induced pluripotent stem cell (ipsc) generation. Cell Transplant 25, 1025–1042. doi:10.3727/096368915X689703

Liu, S., Yang, W., Li, Y. et al. (2023). Fetal bovine serum, an important factor affecting the reproducibility of cell experiments. Sci Rep 13, 1942. doi:10.1038/s41598-023-29060-7

Luengen, A. E., Kniebs, C., Buhl, E. M. et al. (2020). Choosing the right differentiation medium to develop mucociliary phenotype of primary nasal epithelial cells in vitro. Sci Rep 10, 6963. doi:10.1038/s41598-020-63922-8

Mao, H., Wang, Y., Yuan, W. et al. (2009). Ciliogenesis in cryopreserved mammalian tracheal epithelial cells cultured at the air-liquid interface. Cryobiology 59, 250–257. doi:10.1016/j.cryobiol.2009.07.012

Nimmerjahn, F. and Ravetch, J. V. (2008). Fcgamma receptors as regulators of immune responses. Nat Rev Immunol 8, 34–47. doi:10.1038/nri2206

O’Boyle, N., Sutherland, E., Berry, C. C. et al. (2017). Temporal dynamics of ovine airway epithelial cell differentiation at an air-liquid interface. PLoS One 12, e0181583. doi:10.1371/journal.pone.0181583

O’Boyle, N., Sutherland, E., Berry, C. C. et al. (2018). Optimisation of growth conditions for ovine airway epithelial cell differentiation at an air-liquid interface. PLoS One 13, e0193998. doi:10.1371/journal.pone.0193998

Pezzulo, A. A., Starner, T. D., Scheetz, T. E. et al. (2011). The air-liquid interface and use of primary cell cultures are important to recapitulate the transcriptional profile of in vivo airway epithelia. Am J Physiol Lung Cell Mol Physiol 300, L25–31. doi:10.1152/ajplung.00256.2010

Price, P. J. and Gregory, E. A. (1982). Relationship between in vitro growth promotion and biophysical and biochemical properties of the serum supplement. In Vitro 18, 576–584. doi:10.1007/BF02810081

Prytherch, Z., Job, C., Marshall, H. et al. (2011). Tissue-specific stem cell differentiation in an in vitro airway model. Macromol Biosci 11, 1467–1477. doi:10.1002/mabi.201100181

Rauch, C., Feifel, E., Amann, E. M. et al. (2011). Alternatives to the use of fetal bovine serum: Human platelet lysates as a serum substitute in cell culture media. ALTEX 28, 305–316. doi:10.14573/altex.2011.4.305

Redman, E., Fierville, M., Cavard, A. et al. (2024). Cell culture differentiation and proliferation conditions influence the in vitro regeneration of the human airway epithelium. Am J Respir Cell Mol Biol 71, 267–281. doi:10.1165/rcmb.2023-0356MA

Roth, D., Sahin, A. T., Ling, F. et al. (2025). Structure and function relationships of mucociliary clearance in human and rat airways. Nat Commun 16, 2446. doi:10.1038/s41467-025-57667-z

Russell, W. M. S. and Burch, R. L. (1959). The principles of humane experimental technique. Vol. UK, reprinted 1992: Universities Federation for Animal Welfare, Wheathampstaed.

Schaaf, D., Dresen, M., Weldearegay, Y. B. et al. (2026). Mono- and co-infections of primary porcine respiratory cells with bordetella bronchiseptica and streptococcus suis are not affected by the dermonecrotic toxin. Infect Immun e0036625. doi:10.1128/iai.00366-25

Schallmoser, K. and Strunk, D. (2013). Generation of a pool of human platelet lysate and efficient use in cell culture. Methods Mol Biol 946, 349–362. doi:10.1007/978-1-62703-128-8_22

Stival, A. C. S., da Silva, A. C. G. and Valadares, M. C. (2025). Qualitative and quantitative evaluation of fetal bovine serum composition: Toward ethical and best quality in vitro science. NAM J 1, 100047. doi:10.1016/j.namjnl.2025.100047

Tate, J. and Ward, G. (2004). Interferences in immunoassay. Clin Biochem Rev 25, 105–120.

Tu, C., Rudnick, P. A., Martinez, M. Y. et al. (2010). Depletion of abundant plasma proteins and limitations of plasma proteomics. J Proteome Res 9, 4982–4991. doi:10.1021/pr100646w

Umeno, Y., Okuda, A. and Kimura, G. (1989). Proliferative behaviour of fibroblasts in plasma-rich culture medium. J Cell Sci 94 (Pt 3), 567–575. doi:10.1242/jcs.94.3.567

van der Valk, J., Mellor, D., Brands, R. et al. (2004). The humane collection of fetal bovine serum and possibilities for serum-free cell and tissue culture. Toxicol In Vitro 18, 1–12. doi:10.1016/j.tiv.2003.08.009

van der Valk, J., Brunner, D., De Smet, K. et al. (2010). Optimization of chemically defined cell culture media--replacing fetal bovine serum in mammalian in vitro methods. Toxicol In Vitro 24, 1053–1063. doi:10.1016/j.tiv.2010.03.016

van der Valk, J., Bieback, K., Buta, C. et al. (2018). Fetal bovine serum (fbs): Past - present - future. ALTEX 35, 99–118. doi:10.14573/altex.1705101

van Vijven, M., Wunderli, S. L., Ito, K. et al. (2021). Serum deprivation limits loss and promotes recovery of tenogenic phenotype in tendon cell culture systems. J Orthop Res 39, 1561–1571. doi:10.1002/jor.24761

Weber, T., Malakpour-Permlid, A., Chary, A. et al. (2025). Fetal bovine serum: How to leave it behind in the pursuit of more reliable science. Front Toxicol 7, 1612903. doi:10.3389/ftox.2025.1612903

Weldearegay, Y. B., Brogaard, L., Rautenschlein, S. et al. (2025). Primary cell culture systems to investigate host-pathogen interactions in bacterial respiratory tract infections of livestock. Front Cell Infect Microbiol 15, 1565513. doi:10.3389/fcimb.2025.1565513

Wessman, S. J. and Levings, R. L. (1999). Benefits and risks due to animal serum used in cell culture production. Dev Biol Stand 99, 3–8.

Whitcutt, M. J., Adler, K. B. and Wu, R. (1988). A biphasic chamber system for maintaining polarity of differentiation of cultured respiratory tract epithelial cells. In Vitro Cell Dev Biol 24, 420–428. doi:10.1007/bf02628493

Yoon, J. H., Gray, T., Guzman, K. et al. (1997). Regulation of the secretory phenotype of human airway epithelium by retinoic acid, triiodothyronine, and extracellular matrix. Am J Respir Cell Mol Biol 16, 724–731. doi:10.1165/ajrcmb.16.6.9191474

Zhang, P., Cao, L., Ma, Y. Y. et al. (2022). Metagenomic analysis reveals presence of different animal viruses in commercial fetal bovine serum and trypsin. Zool Res 43, 756–766. doi:10.24272/j.issn.2095-8137.2022.093

Zheng, X., Baker, H., Hancock, W. S. et al. (2006). Proteomic analysis for the assessment of different lots of fetal bovine serum as a raw material for cell culture. Part iv. Application of proteomics to the manufacture of biological drugs. Biotechnol Prog 22, 1294–1300. doi:10.1021/bp060121o

