## Supplementary Figures for "Defining a New Standard: Human Platelet Lysate Supports Proliferation and Differentiation of Primary Respiratory Epithelial Cells"

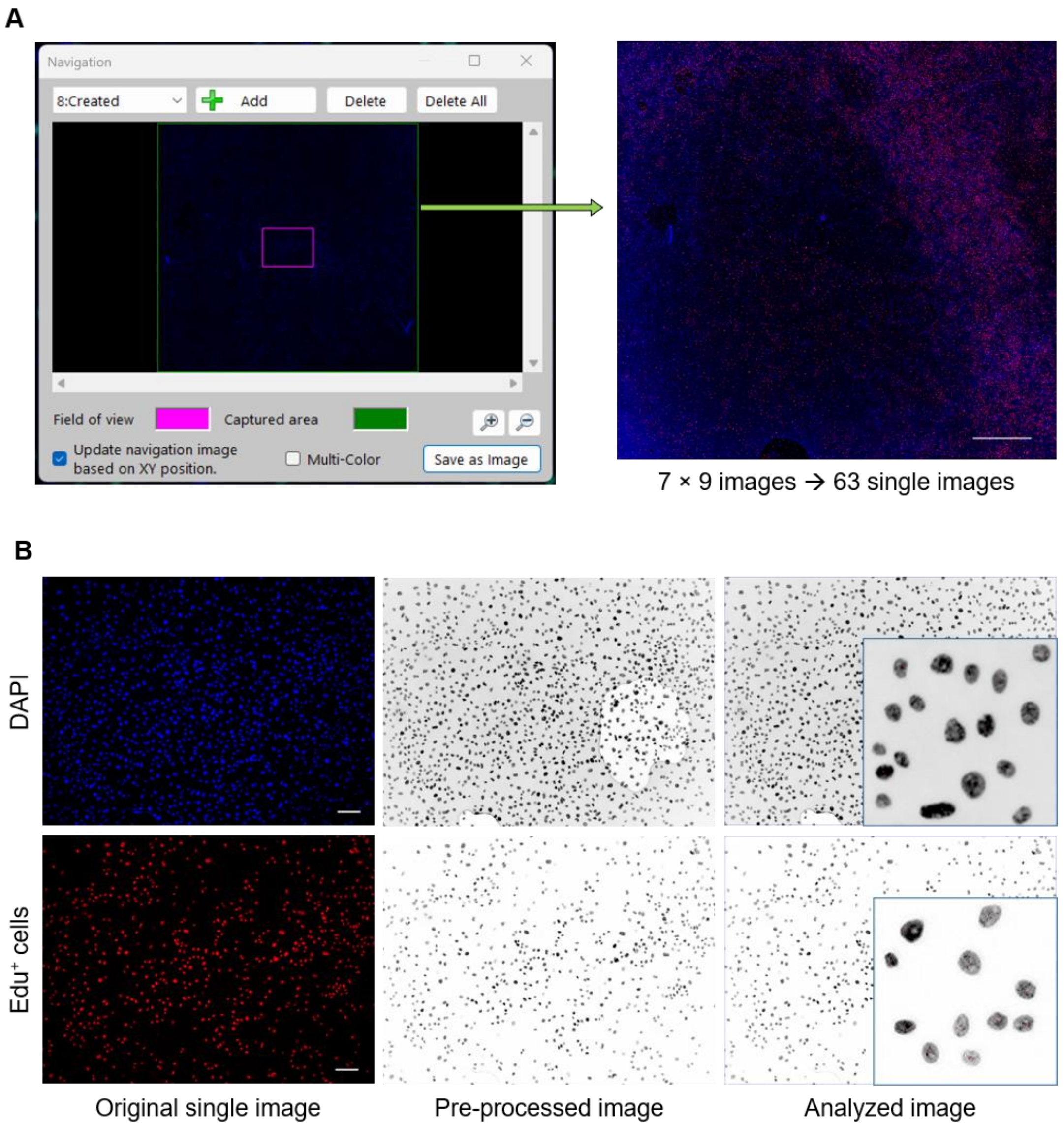


**Fig. S1. Counting of proliferating cells using the ImageJ plug-in Image-based Tool for Counting Nuclei (ITCN).** Proliferating porcine respiratory epithelial cells (PRECs) were labeled using the ClickTech Sensitive EdU Cell Proliferation Kit for Imaging and detected by fluorescence microscopy (EdU$^+$ cells; green). Additionally, nuclei of all cells were labeled with DAPI (blue). (A) The navigation window shows the captured area of each cover slip, consisting of 63 single images. Individual image tiles were captured using the stitching acquisition mode of the Keyence BZ-X800E and assembled into composite images using BZ-X800 Analyzer software. Bar represents 1 mm. (B) Single images were pre-processed prior to analysis with the Image J plug-in ITCN by converting the image to 8-bit and inverting bright and dark signals using ImageJ software. Brightness and contrast were automatically adjusted by the software. Subsequently, pre-processed single images were analyzed using Batch ITCN (Edge: 0 pixels; Nuclei diameter: 30 pixels; Minimum distance: 15 pixels; Threshold: 0.2 for DAPI and 0.1 for EdU$^+$ cells, respectively), labeling each counted cell with a small red dot (enlarged inlay in analyzed images). Bars represent 100 µm.

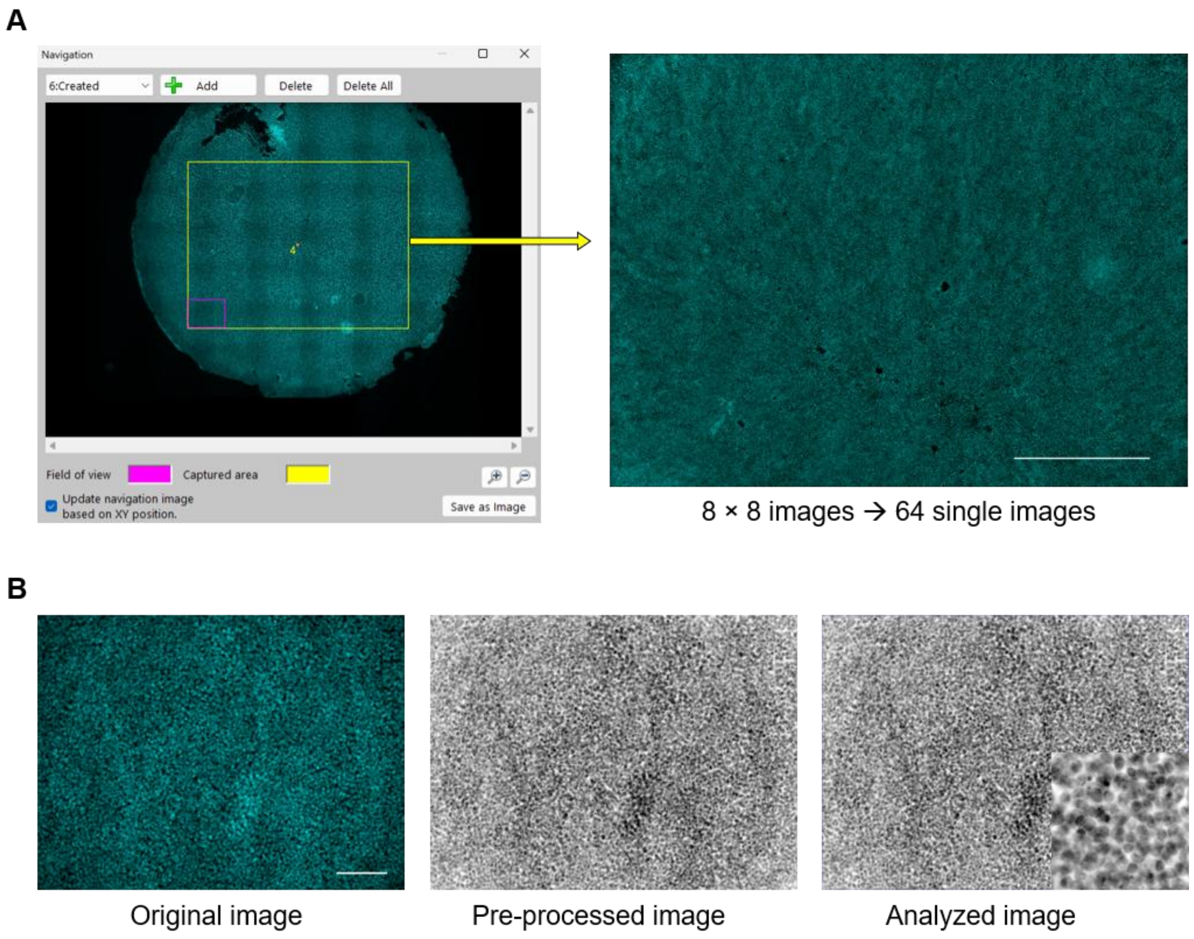


**Fig. S2. Counting cells on transwell filters using the ImageJ plug-in ITCN.** Nuclei were labeled with DAPI (aqua) and visualized using fluorescence microscopy. (A) The navigation window shows the captured area of each transwell filter, consisting of 64 single images. Individual image tiles were captured using the stitching acquisition mode of the Keyence BZ-X800E and assembled into composite images using BZ-X800 Analyzer software. Bar represents 1 mm. (B) Single images were pre-processed prior to analysis with the Image J plug-in ITCN by converting the image to 8-bit and inverting bright and dark signals using ImageJ software. Brightness and contrast were automatically adjusted by the software. Subsequently, pre-processed single images were analyzed using Batch ITCN (Edge: 0 pixels; Nuclei diameter: 20 pixels; Minimum distance: 10 pixels; Threshold: 0.1), labeling each counted cell with a small red dot (enlarged inlay in analyzed images). Bar represents 100 µm.

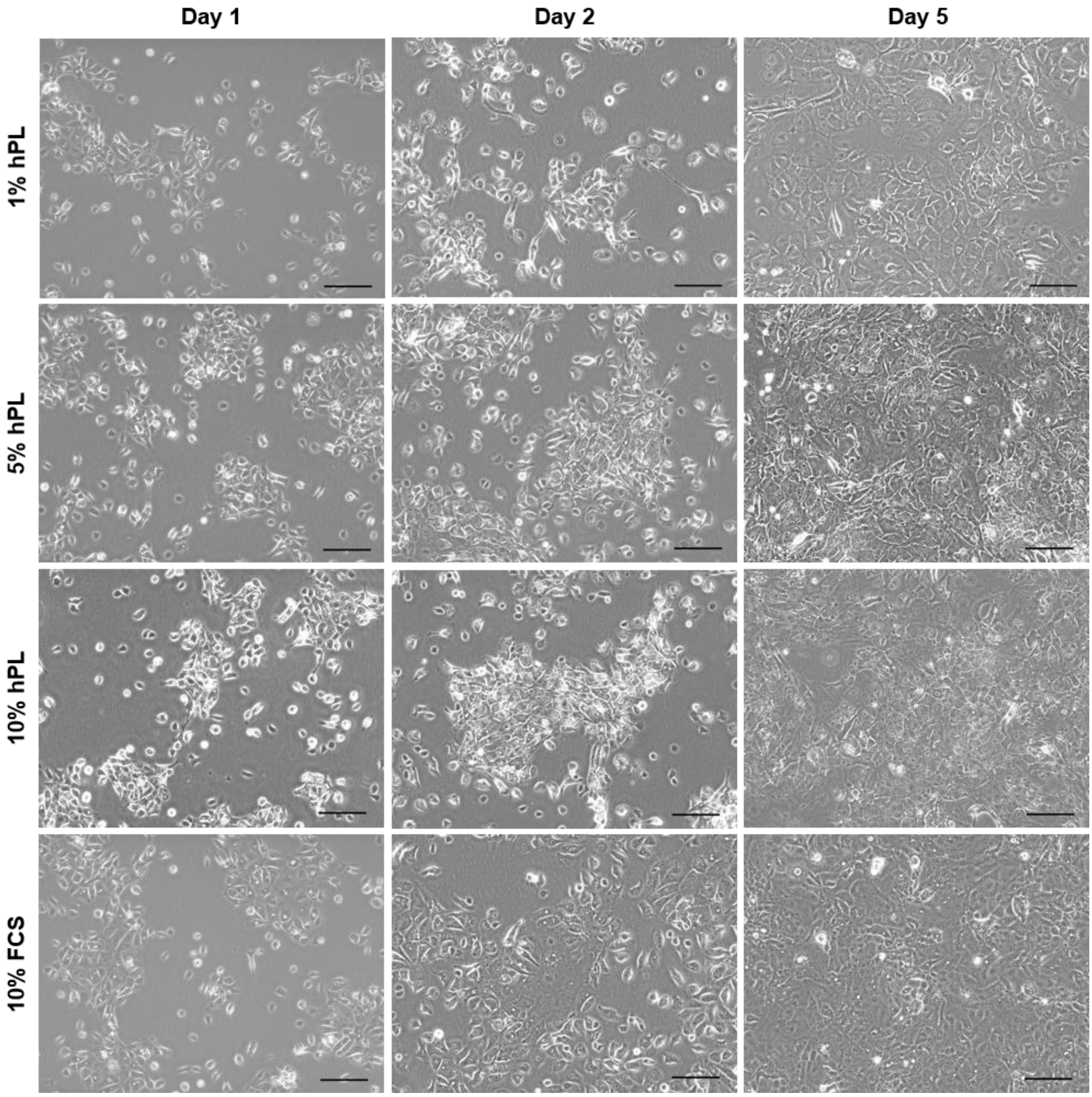


**Fig. S3. Cell morphology of proliferating PRECs.** PRECs were cultivated in Airway Epithelial Cell Growth Medium (AEGM) supplemented with either 1%, 5%, or 10% human platelet lysate (hPL) or 10% fetal calf serum (FCS) for up to five days. Cell morphology was monitored by phase contrast microscopy. Representative images of cells cultivated for one, two, and five days under submerged conditions are shown. Bars represent 100 µm.

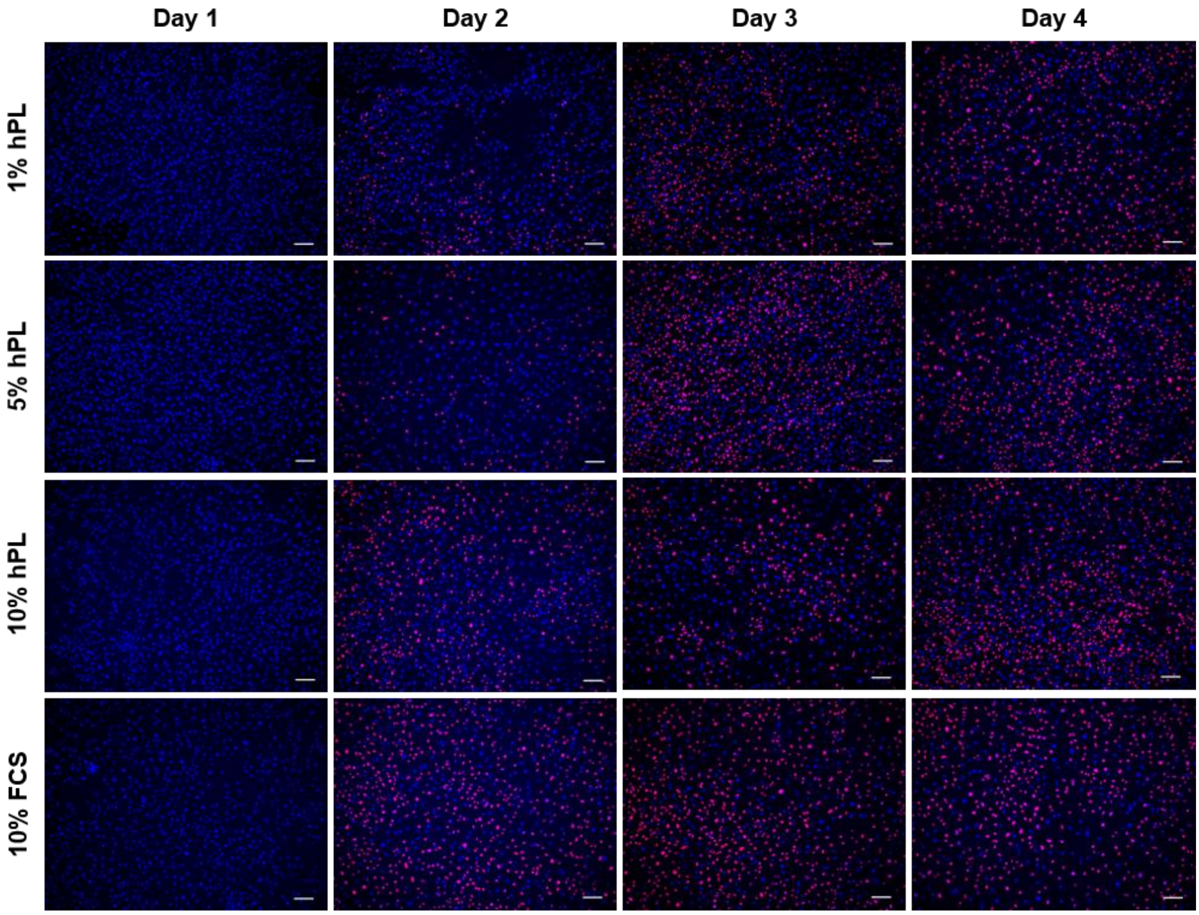


**Fig. S4. Fluorescence microscopy of proliferating PRECs.** Frozen PRECs were seeded at a density of 2 × $10^5$ cells/well on glass cover slips in a 24-well plate. AEGM was supplemented with either 1%, 5%, or 10% hPL or 10% FCS. After one day, medium was changed and 5 µM 5-ethynyl-2'-deoxyuridine (EdU) were added to the cells to label proliferating PRECs. Proliferating ($EdU^+$) cells were then detected by fluorescence microscopy. Representative images of nuclei labeled with 4',6-diamidine-2'-phenylindole (DAPI; blue) and proliferating cells labeled with EdU (red). Bars represent 100 µm.

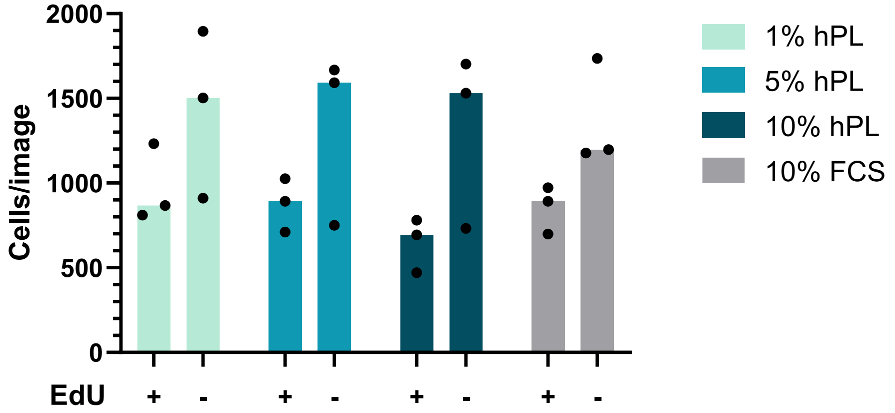


**Fig. S5. Impact of EdU on cell numbers.** PRECs were cultivated in AEGM supplemented with either 1%, 5%, or 10% hPL or 10% FCS in the presence or absence of EdU, respectively, for four days. The number of cells per image was determined by counting nuclei stained with DAPI using the ImageJ plug-in ITCN. The median of three independent experiments is shown. Statistical difference between EdU-treated and non-treated PRECs was analyzed with Mann-Whitney test.

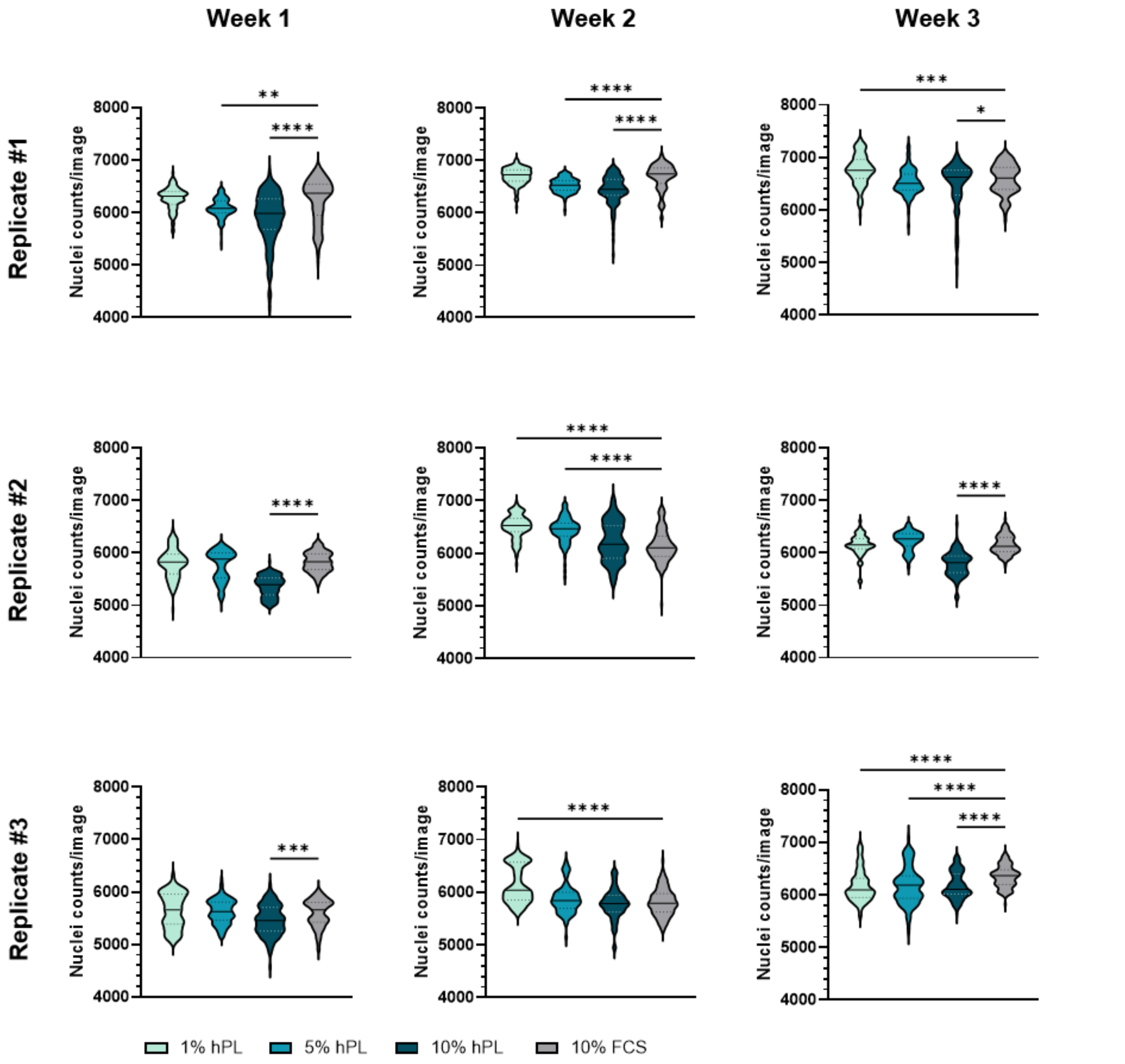


**Fig. S6. Cell numbers per image during differentiation under ALI conditions.** Before PRECs were exposed to ALI conditions, they have been cultivated on transwell filters under submerged conditions in AEGM supplemented with either 1%, 5%, or 10% hPL or 10% FCS for four days. Subsequently, PRECs were differentiated under ALI conditions in medium without any hPL or FCS for up to three weeks. The number of cells per image was determined by counting nuclei labeled with DAPI using the Image J plug-in ITCN. The graphs show nuclei counts per image from two technical replicates each – from at least 60 images, on average from 110-120 images. Violin plots are showing the median as well as the upper and lower quartiles. Significant differences between hPL and 10% FCS were analyzed with one-way ANOVA followed by Tukey's multiple comparisons test (* $p < 0.05$, ** $p < 0.01$, *** $p < 0.001$, **** $p < 0.0001$).

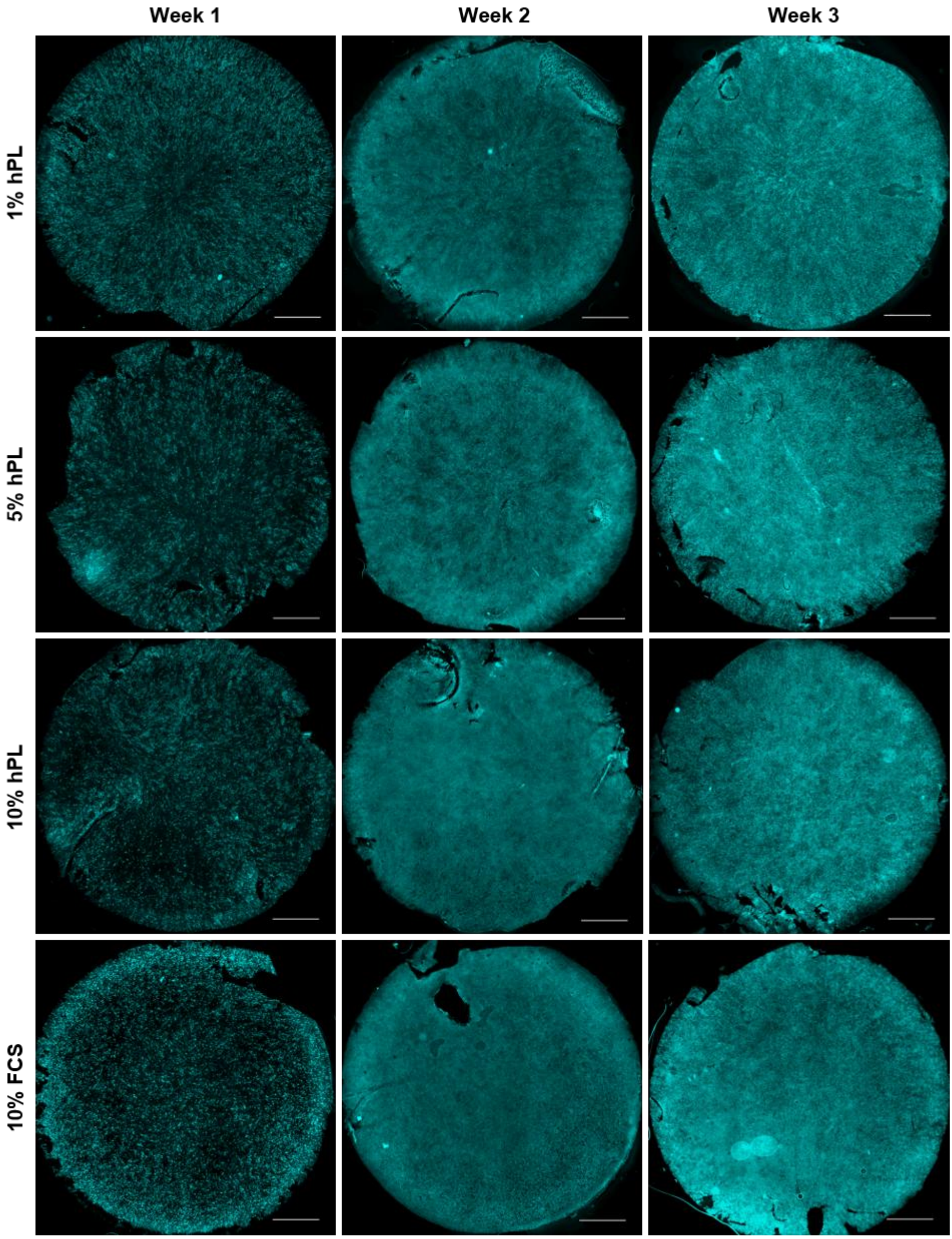


**Fig. S7. Cilia formation of PRECs under air-liquid interface (ALI) conditions.** Before PRECs were exposed to ALI conditions, they have been cultivated on transwell filters under submerged conditions in AEGM supplemented with either 1%, 5%, or 10% hPL or 10% FCS for four days. Subsequently, PRECs were differentiated under ALI conditions in medium without any hPL or FCS for up to three weeks. Immunofluorescence staining of cilia (β-tubulin) was used as an indicator of the level of differentiation under ALI conditions. Composite images of the entire membrane were generated from individual tiles acquired with the Keyence BZ-X800E and processed using BZ-X800 Analyzer. Bars represent 1 mm.
