## Supplementary Table 1 for "Defining a New Standard: Human Platelet Lysate Supports Proliferation and Differentiation of Primary Respiratory Epithelial Cells"

### Defining a New Standard: Human Platelet Lysate Supports Proliferation and Differentiation of Primary Respiratory Epithelial Cells

Anna Richter[1], Jeannine Biermann[1], Marcus Fulde[1], and Désirée Schaaf[1]

[1] *Institute of Microbiology, University of Veterinary Medicine Hannover, Hannover, Germany*

### Cell Culture Media

| Additive | Company | Cat. No. | Final concentration |
|---|---|---|---|
| **Wash Medium** | | | |
| DMEM (high glucose, with pyruvate) | Thermo Fisher Scientific | 41966052 | |
| Penicillin/Streptomycin | Merck | P4333 | 100 U, 0.1 mg/ml |
| Amphotericin B | Merck | A2942 | 2.5 µg/ml |
| Gentamicin | Carl Roth | HN09.2 | 50 µg/ml |
| **Incubation Medium** | | | |
| DMEM (high glucose, with pyruvate) | Thermo Fisher Scientific | 41966052 | |
| Protease | Merck | P5147 | 1 mg/ml |
| DNase I | Merck | 11284932001 | 10 µg/ml |
| Penicillin/Streptomycin | Merck | P4333 | 100 U, 0.1 mg/ml |
| Amphotericin B | Merck | A2942 | 2.5 µg/ml |
| Gentamicin | Carl Roth | HN09.2 | 50 µg/ml |
| **Protease Stopping Buffer** | | | |
| Phosphate Buffered Saline (PBS) | Merck | D8537 | |
| Bovine Serum Albumin (BSA) | Carl Roth | 8076.3 | 0.5% |
| Ethylenediaminetetraacetic Acid (EDTA) | Carl Roth | 8043.1 | 2 mM |
| **Airway Epithelial Cell Growth Medium (AEGM)** | | | |
| Airway Epithelial Cell Basal Medium | PromoCell | C-21260 | |
| Airway Epithelial Cell Growth Medium SupplementMix | PromoCell | C-39165 | |
| Retinoic Acid | Merck | R2625 | 15 ng/ml |
| Y-27632 Dihydrochloride | Tocris | 1254 | 10 µM |
| Fetal Bovine/Calf Serum (FBS/FCS)<br>**OR**<br>Human Platelet Lysate (hPL) | Bio&SELL<br><br>PL BioScience | FBS.S.0615<br><br>PE30611 | 10%<br><br>1%/ 5%/10% |
| Penicillin/Streptomycin | Merck | P4333 | 100 U, 0.1 mg/ml |
| Amphotericin B | Merck | A2942 | 2.5 µg/ml |
| Gentamicin | Carl Roth | HN09.2 | 50 µg/ml |
| **ALI Medium** | | | |
| Airway Epithelial Cell Basal Medium | PromoCell | C-21260 | Mixture 1:1 |
| DMEM (high glucose, with pyruvate) | Thermo Fisher Scientific | 41966052 | |
| Airway Epithelial Cell Growth Medium SupplementMix | PromoCell | C-39165 | |
| Bovine Serum Albumin | Merck | A7638 | 500 µg/ml |

| | | | |
|---|---|---|---|
| Retinoic Acid | Merck | R2625 | 15 ng/ml |
| Penicillin/Streptomycin | Merck | P4333 | 100 U, 0.1 mg/ml |
| **Freezing Medium** | | | |
| AEGM | | | |
| FCS | Bio&SELL | FBS.S.0615 | 30% |
| Dimethyl sulfoxide (DMSO) | Merck | D2650 | 10% |

### Other Reagents

| Reagent | Company | Cat. No. | Final concentration |
|---|---|---|---|
| Collagen I | Merck | C3867 | 10 µg/cm$^2$ |
| Collagen IV | Merck | C7521 | 7.5 µg/cm$^2$ |
| Trypsin-EDTA (TE) | Thermo Fisher Scientific | 15400054 | 0.05% |
| Formaldehyde | Carl Roth | CP10.1 | 3.7% |
| Saponin | Carl Roth | 4185.1 | 1% |
| BSA | Carl Roth | 8076.3 | 3% |

### Buffer and Reagents for Immunofluorescence Staining

| Reagent | Company | Cat. No. | Final concentration |
|---|---|---|---|
| Anti-β-Tubulin Antibody (clone TUB 2.1, CY3 conjugate) | Merck | C4585 | 1:500 |
| 4', 6-diamidino-2-phenylindole dihydrochloride (DAPI) | Cell Signaling | 4083S | 0.5 µg/ml |
| ProLong Gold Antifade Reagent | Cell Signaling | 9071S | |
| **Blocking Buffer** | | | |
| PBS | | | |
| Glycine | Carl Roth | 3908.3 | 0.1 M |
| Goat Serum | Merck | S6898 | 5% |
| Tergitol™ 15-S-9 | Carl Roth | 9975.1 | 0.5% |
| Tween® 20 | Carl Rith | 9127.2 | 0.05% |
| **Antibody Dilution Buffer** | | | |
| PBS | | | |
| BSA | Carl Roth | 8076.3 | 1% |
| Tergitol™ 15-S-9 | Carl Roth | 9975.1 | 0.5% |
| Tween® 20 | Carl Rith | 9127.2 | 0.05% |
